# Eldecalcitol is more effective for promoting osteogenesis than alfacalcidol in *Cyp27b1*-knockout Mice

**DOI:** 10.1101/349837

**Authors:** Yoshihisa Hirota, Kimie Nakagawa, Keigo Isomoto, Toshiyuki Sakaki, Noboru Kubodera, Maya Kamao, Naomi Osakabe, Yoshitomo Suhara, Toshio Okano

## Abstract

Calcium (Ca) absorption from the intestinal tract is promoted by active vitamin D (1α,25D_3_). Vitamin D not only promotes Ca homeostasis, but it also inhibits bone resorption and promotes osteogenesis, thus playing a role in the maintenance of normal bone metabolism. Because 1α,25D_3_ plays an important role in osteogenesis, vitamin D formulations, such as alfacalcidol (ALF) and eldecalcitol (ELD), are used for treating osteoporosis. While it is known that, in contrast to ALF, ELD is an active ligand that directly acts on bone, the reason for its superior osteogenesis effects is unknown. *Cyp27b1*-knockout mice (*Cyp27b1*^−/−^ mice) are congenitally deficient in 1α,25D_3_ and exhibit marked hypocalcemia and high parathyroid hormone levels, resulting in osteodystrophy involving bone hypocalcification and growth plate cartilage hypertrophy. However, because the vitamin D receptor is expressed normally in *Cyp27b1*^−/−^ mice, they respond normally to 1α,25D_3_. Accordingly, in *Cyp27b1*^−/−^ mice, the pharmacological effects of exogenously administered active vitamin D derivatives can be analyzed without being affected by 1α,25D_3_. We used *Cyp27b1*^−/−^ mice to characterize and clarify the superior osteogenic effects of ELD on the bone in comparison with ALF. The results indicated that compared to ALF, ELD strongly induces ECaC2, calbindin-D_9k_, and CYP24A1 in the duodenum, promoting Ca absorption and decreasing the plasma concentration of 1α,25D_3_, resulting in improved osteogenesis. Because bone morphological measurements demonstrated that ELD has stronger effects on bone calcification, trabecular formation, and cancellous bone density than ALF, ELD appears to be a more effective therapeutic agent for treating postmenopausal osteoporosis, in which cancellous bone density decreases markedly. By using *Cyp27b1*^−/−^ mice, this study was the first to succeed in clarifying the osteogenic effect of ELD without any influence of endogenous 1α,25D_3_. Furthermore, ELD more strongly enhanced bone mineralization, trabecular proliferation, and cancellous bone density than did ALF. Thus, ELD is expected to show an effect on postmenopausal osteoporosis, in which cancellous bone mineral density decreases markedly.

## Introduction

Vitamin D_3_, which is incorporated into the body through diet and synthesis in the skin, undergoes 25-hydroxylation by vitamin D 25-hydroxylase (CYP2R1 and CYP27A1) in the liver to become 25-hydroxyvitamin D3 (25D3). Next, 25-hydroxyvitamin D 1α-hydroxylase (CYP27B1) in the kidneys catalyzes 1α-hydroxylation to form 1α,25D_3_. Meanwhile, 25D_3_ undergoes 24-hydroxylation by 25-hydroxyvitamin D 24-hydroxylase (CYP24A1) to be metabolized into 24,25-dihydroxyvitamin D_3_ (24,25D_3_).

In living organisms, 1α,25D_3_ is transported in the plasma when bound with vitamin D-binding protein (DBP) to reach target tissues [1], such as the bones, kidneys, parathyroid glands and small intestine, where it binds to the vitamin D receptor (VDR), which belongs to the intranuclear steroid hormone receptor super family[2]. VDR, when bound to 1α,25D_3_, forms a dimer with retinoid X receptor before binding with vitamin D responsive element (VDRE) in the target gene promoter to mutually interact with various transcriptional coactivators and basal transcription factors to activate promoters inducing target gene expression[3–5]. Thus, 1α,25D_3_ plays important roles in balancing Ca metabolism in living organisms by modulating target gene expression and promoting active Ca absorption in the intestinal tract as well as Ca reabsorption in the kidneys, osteogenesis, and bone resorption [6]. The plasma 1α,25D_3_ concentration is normally maintained in the range of 20–70 pg/mL by CYP27B1. Plasma Ca, P, parathyroid hormone (PTH) or calcitonin level induce CYP27B1 expression in the kidneys, increasing 1α,25D_3_ production [7]. In turn, 1α,25D_3_ inhibits VDR-mediated CYP27B1 expression at the transcription level to create a negative feedback loop [8]. Thus, the production volume of 1α,25D_3_ is strictly controlled.

Not only does 1α,25D_3_ promote active Ca absorption from the intestinal tract, it also inhibits PTH secretion [9], suppresses bone resorption, and promotes osteogenesis in order to maintain normal bone remodeling and bone quantity. However, as 1α,25D_3_ strongly promotes intestinal Ca resorption, when administered in pharmacological quantities, there is a risk of hypercalcemia [10]. Therefore, alfacalcidol (ALF), a prodrug of 1α,25D_3_, is mainly used clinically and has long been the most widely used prodrug for this purpose in Japan. Because ALF is metabolized in the liver to 1α,25D_3_ via 25-hydroxylation, it has been used clinically as a therapeutic agent for treating diseases involving abnormal vitamin D metabolism, such as chronic kidney failure and vitamin D-dependent rickets. In addition, it has become widely applied as a vitamin D formulation for osteoporosis. ALF reportedly increases bone density, inhibits bone fractures, and suppresses the breakdown of the trabecular structure [11–13].

In recent years, vitamin D formulations improving bone metabolism have been developed for the treatment of osteoporosis. Eldecalcitol (ELD) was discovered through *in-vivo* screening using ovariectomized rats (OVX rats) as an osteoporosis model animal. ELD is a compound with a functional group introduced at 2β of 1α,25D_3_ [14]. Unlike ALF, ELD is not need to be metabolized; it is an active vitamin D formulation in which the compound itself acts as an active ligand [15]. ELD exhibits lower “binding affinity” for VDR than 1α,25D_3_, but stronger binding affinity to DBP [16,17]. As this strong binding affinity to DBP increases stability in the plasma, it features a long plasma half-life [17,18]. Its PTH secretion-inhibitory effects reportedly are weaker than those of 1α,25D_3_ [19].

In elderly rats and OVX rats, ELD increases thoracic vertebra bone density and bone strength [18]. Further, ELD increases bone quantity and osteogenesis speed in normal and OVX rats [20]. These studies using osteoporotic animals have shown that ELD offers superior bone quantity-increasing effects. Clinical experiments have indicated that ELD inhibits bone resorption in a dosage-dependent manner without affecting osteogenesis, thereby increasing thoracic vertebra bone density [21]. However, the mechanism underlying the superior osteogenic effects of ELD remains unclear. In particular, because it has been reported that the plasma 1α,25D_3_ concentration decreases upon ELD administration, it is unknown whether its superior osteogenic effects are direct effects or are mediated by decreased plasma 1α,25D_3_ [22].

*Cyp27b1*^−/−^ mice have been bred by multiple research teams and are used in studies of the physiological functions of CYP27B1 [23–26]. *Cyp27b1*^−/−^ mice exhibit marked hypocalcemia and high PTH levels after weaning. This results in decreased growth, with osteodystrophy involving bone hypocalcification and growth plate cartilage hypertrophy [23–26]. In *Cyp27b1*^−/−^ mice, although VDR is expressed normally, 1α,25D_3_ deficiency occurs because 1α-hydroxylation of 25D_3_ by CYP27B1 does not occur. Accordingly, *Cyp27b1*^−/−^ mice are an effective animal model for analyzing the physiological actions of exogenously administered active vitamin D derivatives in an environment without endogenous 1α,25D_3_. Therefore, to clarify the osteogenic effects of ELD in an environment not affected by endogenic 1α,25D_3_, we used *Cyp27b1*^−/−^ mice in this study. In addition, we investigated the characteristics of ELD osteogenesis in comparison with ALF, which is clinically applied as a therapeutic agent for osteoporosis.

## Materials and methods

### Materials

Heterozygous *Cyp27b1* knockout mice (*Cyp27b1*^+/−^ mice) were produced according to previously reported methods [26]. Male and female *Cyp27b1*^+/−^ mice were bred to create *Cyp27b1*^+/+^, *Cyp27b1*^+/−^, and *Cyp27b1*^−/−^ mice. The mice were fed Diet 11 [CLEA Japan, Suita, Japan], a vitamin D-deficient feed, supplemented with 2.4 IU/g of vitamin D_3_ (Diet11+D). The feed was manufactured by CLEA Japan at our request. Medium-chain triglycerides, 1α-hydroxyvitamin D_3_ (ALF), and 2β-hydroxypropoxy-1α,25-dihydroxyvitamm D_3_ (ELD) were supplied by Chugai Pharmaceutical Co., Ltd. (Tokyo, Japan). The ATDC5 cells used in this study are cartilage cells isolated from mouse teratocarcinoma (RIKEN Cell Bank, Tsukuba, Japan). Organic solvents of HPLC grade were purchased from Nacalai Tesque (Kyoto, Japan). Other chemicals were highest-grade commercial chemicals.

### Ethics statement

All cell experimental protocols were performed in accordance with the Guidelines for Cell Experiments at Kobe Pharmaceutical University and were approved by The Ethics Committee of Kobe Pharmaceutical University, Kobe Japan. All animal experimental protocols were performed in accordance with the Guidelines for Animal Experiments at Kobe Pharmaceutical University and were approved by The Animal Research and Ethics Committee of Kobe Pharmaceutical University, Kobe Japan. All surgery was performed under sodium pentobarbital anesthesia, and all efforts were made to minimize suffering.

### Rearing conditions and body-weight measurements

All mice were weaned at 3 weeks of age. After weaning, they were allowed to freely feed on a diet of solid food (Diet 11+D) and deionized water. The mice were raised in an enclosure with controlled temperature and humidity (23 ± 1°C, 60 ± 1%). The mice were raised until 9 weeks of age (administration period: 6 weeks). Body weight was measured once per week starting directly after weaning (3, 4, 5, 6, 7, 8 and 9 weeks of age).

### ALF and ELD dosing solutions

ALF and ELD dosing solutions were formulated by dissolving 100 μL of ethanol containing 10 μg/mL of ALF or ELD into 7.9 mL of a medium-chain triglyceride solution. The vehicle dosing solution was prepared similarly using 100 μL of ethanol solution. The solutions were administered orally 3 times per week to achieve a dosage of 0.25 μg/kg body weight.

### Plasma Ca and PTH concentration measurements

Plasma was collected weekly in heparin-coated capillary tubes (Mylar^®^ Wrapped 75MM Hematocrit Tubes; Drummond Scientific Company, Pennsylvania, US) from the caudal vein, under anesthesia. The plasma Ca concentration was measured using a Ca measurement kit (Wako Calcium C-Test; FUJIFILM Wako Pure Chemical, Osaka, Japan). At 9 weeks of age, cardiac plasma was collected and serum was separated by centrifugation. The plasma PTH concentration was measured using a mouse PTH measurement kit (Mouse Intact PTH ELISA Kit; Immutopics, CA, USA).

### Reverse-transcription quantitative PCR

Total RNA was isolated from the mouse duodenum and liver using Isogen (Nippon Gene, Tokyo, Japan), in accordance with the manufacturer’s protocol. First-strand cDNA was generated using AMV reverse transcriptase (Takara Bio, Kusatsu, Japan). PCRs were conducted using a SYBR Green Core Reagent Kit (PE Biosystems, Foster City, CA, USA) on a CFX96 Real-time PCR System (Bio-Rad, Hercules, CA, USA), according to the manufacturer’s protocol. Primers were designed to target mouse epithelial calcium channel 2 (ECaC2) (GenBank accession number NM_022413; forward primer, base pairs 1433–1452; reverse primer, 1859–1878), mouse calbindin-D_9_k (NM_004057.2, forward primer, 111–130; reverse primer, 227–246), mouse Cyp2r1 (1.XM_006507838.3, forward primer, 1262-1281; reverse primer, 1345-1364), mouse Cyp24a1 (AK159527.1, forward primer, 738-757; reverse primer, 839-858), and mouse β-actin (X03672; forward primer, 250–271; reverse primer, 305–326) as a control. Primer specificity was evaluated by electrophoresis of the PCR product.

### Soft X-ray imaging of the femur and tibia

Mice were anesthetized and then euthanized by means of cardiac plasma collection. The femurs and tibias were collected. SOFTEX soft X-ray imaging equipment (CMB-2; SOFTEX, Kanagawa, Japan) was used to acquire soft X-ray images of the femurs and tibias.

### Femur and bone morphological measurements

Parameters related to bone structure, bone shape, and bone resorption were analyzed in non-decalcified specimens of femurs obtained from female mice after 6 weeks of administrations. Total bone density, cortical bone density, and cancellous bone density of the femoral metaphysis, as well as cortical bone density and cortical bone thickness in the diaphysis were measured by peripheral quantitative computed tomography (pQCT). Horizontal, vertical and twisting bone strength in the femur were measured by pQCT. Two-dimensional (2D) and three-dimensional (3D) structural analyses of femoral trabeculae were conducted by micro-CT. The pQCT and micro CT analyses were performed at Kureha Special Laboratory (Fukushima, Japan).

### Bone histological staining

Bone-tissue sections were prepared by Kureha Special Laboratory. Bone calcification was assessed by von Kossa staining, cartilage formation by toluidine blue staining, and osteoid formation by Villanueva staining of the bone specimens. The bone sections were observed by bright-field microscopy using an all-in-one fluorescence microscope (BZ-8000; KEYENCE, Osaka, Japan). To analyze bone formation speed by calcein double labeling, mice were administered 0.1 mL of calcein dosing solution per 10 g of body weight 4 days prior to and 1 day prior to tissue collection. The specimens were observed under the BZ-8000 microscope using an excitation wavelength of 480 nm and emission wavelength of 505 nm.

### ATDC5 cell culture

ATDC5 cells were maintained in Dulbecco’s modified Eagle’s and Ham-F12 [DMEM/F12] (Nakalai Tesque) supplemented with 1% penicillin, 1% streptomycin, and 10% fetal bovine serum (FBS, Gibco/BRL) at 37°C in a humidified atmosphere with 5% CO_2_. The medium was replaced every other day.

### Cell proliferation assay

ATDC5 cells reached confluence 3 days after seeding. At day 5, cells were exposed to chondrogenic medium consisting of DMEM/Ham’s F-12 (1:1) containing 5% FBS, 10 μg/ml human transferrin (Boehringer Mannheim, Mannheim, Germany), and 3 × 10^−8^ M sodium selenite (Sigma Chemical, St. Louis, MO, USA), 10 μg/ml insulin (FUJIFILEM Wako Pure Chemical, Osaka, Japan) in 96-well plates (1 × 10^4^ cells/well). In addition, 10^−7^–10^−10^ M 1α,25D_3_, ALF, ELD, or 10^−6^–10^−9^ M 25D_3_ was added. The media were changed every other day. Proliferation of ATDC5 cells was monitored using the cell proliferation agent WST-1 (Nakalai Tesque). Following addition of WST-1 reagent, the optical density at 450 nm was read.

### Alcian blue staining

ATDC5 cells were plated in 12-multiwell plates and cultured in chondrogenic medium with 10^−7^–10^−10^ M 1α,25D_3_, ALF, ELD, or 10^−6^–10^−9^ M 25D_3_ using the method described above. The cells were rinsed with PBS and fixed in 95% methanol for 20 min, and then stained with 0.1% alcian blue 8GX (FUJIFILEM Wako Pure Chemical) in 0.1 M HCl overnight. The stained cells were rinsed with PBS three times and observed by microscopy.

### Transfection and luciferase activity assays

To asses the activity via the VDRE, ATDC5 cells (2 × 10^5^) were suspended in 2 mL of medium and transfected with 0.5 μg of pGVB2-rat Cyp24a1 luciferase reporter vector (Toyo Ink, Tokyo, Japan) containing a rat *Cyp24a1* gene promoter including two VDREs and 0.25 μg of pRL-CMV vector (Toyo Ink) as an internal control, using Tfx-50 reagent (Promega, Madison, WI). The cells were incubated with 10^−7^–10^−10^ M 1α,25D_3_, ALF, ELD, or 10^−6^–10^−9^ M 25D_3_ for 48 h. Next, the binding capacity to VDR was examined by using the VDR-GAL4 one-hybrid luciferase system. ATDC5 cells (2 × 10^5^) were suspended in 2 mL of medium and transfected with 0.5 μg of pM-human VDR incorporating whole human VDR cDNA upstream of the DNA-binding domain of the *GAL4* gene, which is a transcriptional regulator in yeast, 0.5 μg of pGVP2-GAL2 containing the SV40 promoter incorporating the binding site of rat *GAL4* upstream of the firefly luciferase gene, and 0.25 μg of pRL-CMV vector (pGVB2 vector; Toyo Ink) as an internal control, using Tfx-50 reagent (Promega). The cells were incubated with 10^−9^ M 1α,25D_3_, ALF, ELD, or 10^−7^ M 25D_3_ and 0, 5, or 25 μg/mL DBP for 48 h. The luciferase activities of the cell lysates were measured with a luciferase assay kit (Toyo Ink), according to the manufacturer’s instructions. Transactivation measured as luciferase activity was standardized to the luciferase activity of the same cells determined with the Sea Pansy luciferase assay system as a control (Toyo Ink).

### Role of vitamin D derivatives in megalin-knockdown cells

Stealth RNAi *siMegalin* is a 25-base-pair (bp) duplex oligoribonucleotide of the reported mouse megalin mRNA sequence. Alexa Fluor-labeled siRNA (Thermo Fisher Scientific, CA, USA) was used as control siRNA. ATDC5 cells were transfected with 50 pmol of each siRNA with the use of Lipofectamine RNAiMAX (Thermo Fisher Scientific) in 1 mL of Opti-MEM serum-free medium (Thermo Fisher Scientific), in accordance with the manufacturer’s instructions. Megalin mRNA was quantified by real-time PCR as described above. The obtained megalin-knockdown cells transfected with 0.5 μg of pGVB2-rat Cyp24a1 vector and 0.25 μg of pRL-CMV vector using Tfx-50 reagent. The cells were incubated with 10^−9^ M 1α,25D_3_, ALF, ELD, or 10^−7^ M 25D_3_ for 48 h. To assay transcriptional activity, the luciferase assay described above was used.

### Statistical analysis

Data are expressed as the mean ± SEM. The number of specimens is n = 5. Means were compared using an unpaired Student’s *t*-test or Dunnett’s test and Tukey–Kramer’s honestly significant difference (HSD) test. *P* < 0.05 was considered significant.

## Results

### Effects of ALF or ELD administration on body weight changes and calcium metabolism in *Cyp27b1*-knockout mice

We monitored body weight in *Cyp27b1*^+/+^ and *Cyp27b1*^−/−^ mice that were administered ALF, ELD, or vehicle for 6 weeks after weaning. From five weeks of age, *Cyp27b1*^−/−^ mice exhibited significantly lower body weight than the *Cyp27b1*^+/+^ mice. In contrast, in *Cyp27b1*^−/−^ mice administered ALF or ELD, body weight increased to levels similar to those observed in *Cyp27b1*^+/+^ mice (Fig 1A). Although a tendency for hypocalcemia was observed in *Cyp27b1*^−/−^ mice treated with vehicle, the administration of ALF or ELD significantly increased plasma Ca concentration. In the ALF and ELD administration groups, both *Cyp27b1*^+/+^ and *Cyp27b1*^−/−^ mice exhibited normal or slightly high calcium Ca concentration as of 6 weeks of age (Fig 1B). Plasma PTH concentration was markedly higher in *Cyp27b1*^−/−^ vehicle-administered mice than in *Cyp27b1*^+/+^ vehicle-administered mice. In *Cyp27b1*^−/−^ mice administered ALF or ELD, plasma PTH was significantly decreased as compared to vehicle-treated mice, and at levels similar to those noted for *Cyp27b1*^+/+^ mice (Fig 1C). Next, we measured the duodenal mRNA expression of *ECaC2* and *calbindin-D_9_k*, which encode calcium metabolism-modulating proteins. In *Cyp27b1*^+/+^ mice administered ELD, mRNA expression levels of *ECaC2* and *calbindin-D_9_k* in the duodenum are significantly increased as compared to those in vehicle- and ALF-administered mice. In *Cyp27b1*^−/−^ mice administered ALF or ELD, mRNA expression levels of both genes tended to increase (Fig 1D,E). In addition, we measured mRNA expression levels of *CYP2R1* and *CYP27A1,* which encode enzymes involved in vitamin D metabolism in the liver. *CYP2R1* and *CYP27A1* mRNA expression levels did not exhibit any significant changes as a result of ALF or ELD administration (Fig 1F, G). Both *Cyp27b1*^+/+^ and *Cyp27b1*^−/−^ mice administered ALF or ELD exhibited significantly increased *CYP24A1* mRNA expression compared to the respective vehicle-treated groups. Particularly marked increases were noted in the ELD administration groups (Fig 1H).

**Fig. 1.**
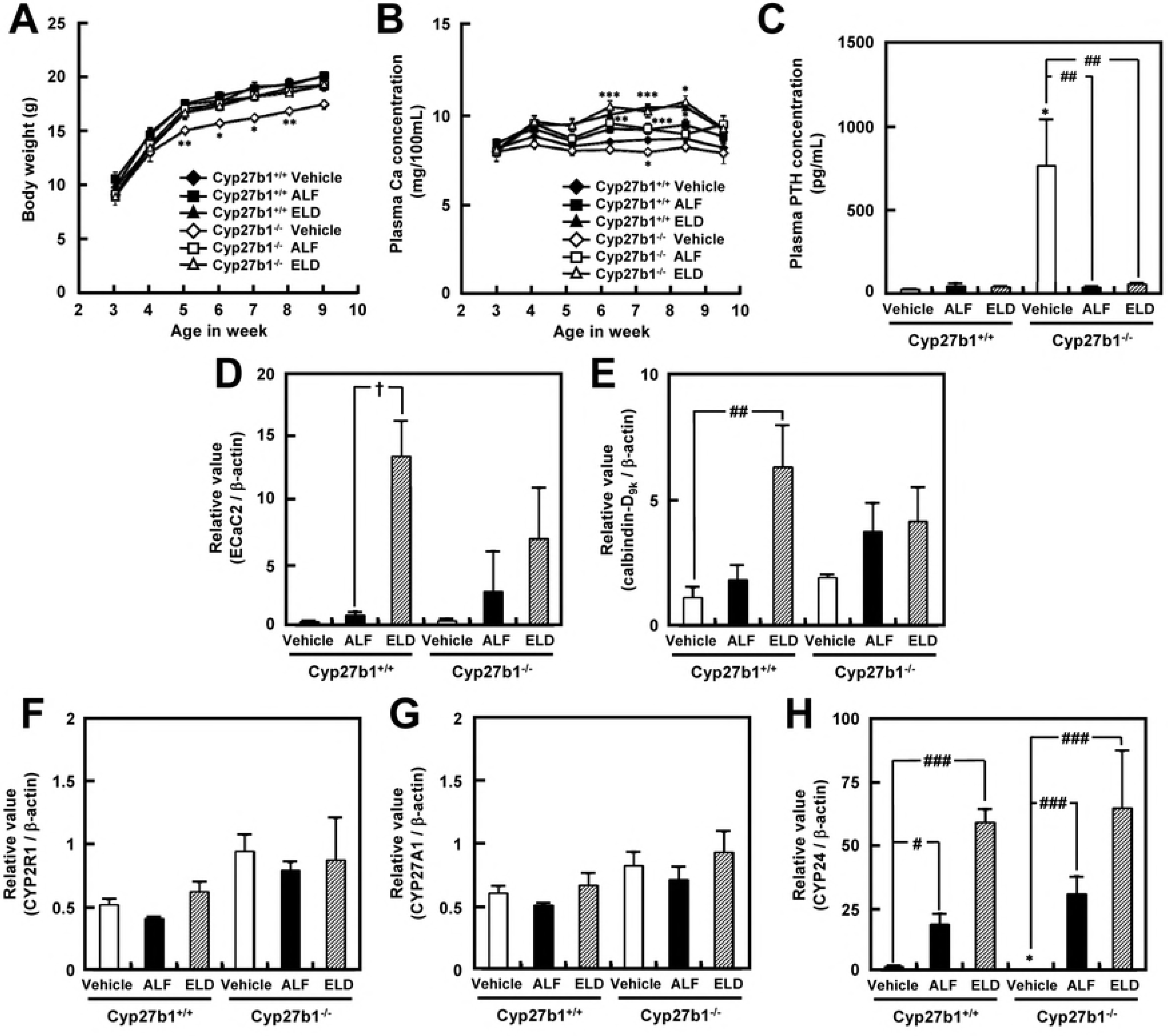
Effects of 6-week ALF or ELD administration in *Cyp27b1*^+/+^ or *Cyp27b1*^−/−^ mice. (A) Body weight. \**P* < 0.05, \*\**P* < 0.01 vs. 3-week old mice, Dunnett’s test. (B) Plasma Ca concentration. Significantly different from each 3-week old mice; \**P* < 0.05, \*\**P* < 0.01, \*\*\**P* < 0.001, Dunnett’s test. (C) Plasma PTH concentration. \**P* < 0.05 vs. *Cyp27b1*^+/+^ mice administered vehicle, Student’s *t*-test. ^##^*P* < 0.01 vs. *Cyp27b1*^−/−^ mice administered vehicle, Dunnett’s test. (D) ECaC2 mRNA expression level in the duodenum. † *P* < 0.05 vs. *Cyp27b1*^+/+^ mice administered ALF, Student’s *t*-test. (E) Calbindin-D_9k_ mRNA expression level in the duodenum. ^##^*P* < 0.01 vs. *Cyp27b1*^+/+^ mice administered vehicle, Student’s *t*-test. (F) CYP2R1 mRNA expression levels in the liver. (G) CYP27A1 mRNA expression levels in the liver. (H) CYP24A1 mRNA expression level in the kidneys. ^#^*P* < 0.05, ^###^*P* < 0.001 vs. *Cyp27b1*^+/+^ or *Cyp27b1*^−/−^ mice administered vehicle, Dunnett’s test.

### External appearance and soft X-ray images of the femurs of *Cyp27b1*-knockout mice administered ALF or ELD

The femurs and tibias of *Cyp27b1*^−/−^ mice administered vehicle were shorter than those of *Cyp27b1*^+/+^ mice administered vehicle, and hypertrophy was noted in the epiphyseal region. In contrast, the external appearance of the bones of *Cyp27b1*^−/−^ mice that were administered ALF or ELD was nearly the same as that of the bones of *Cyp27b1*^+/+^ mice (Fig 2A). Soft X-ray images of femurs and tibias demonstrated that in the *Cyp27b1*^−/−^ mice administered vehicle, the center was black and hollow compared to that in *Cyp27b1*^+/+^ mice, with an unclear shape, indicating hypocalcification. White, clear X-ray images similar to those obtained for *Cyp27b1*^+/+^ mice were observed in the *Cyp27b1*^−/−^ mice administered ALF or ELD, suggesting that the hypocalcification of *Cyp27b1*^−/−^ mice had been improved. In addition, although the femoral bones were significantly shorter in *Cyp27b1*^−/−^ mice administered vehicle than in *Cyp27b1*^+/+^ mice administered vehicle, the bones were longer in *Cyp27b1*^−/−^ mice administered ALF or ELD, with lengths being similar to those in *Cyp27b1*^+/+^ mice (Fig 2B).

**Fig. 2.**
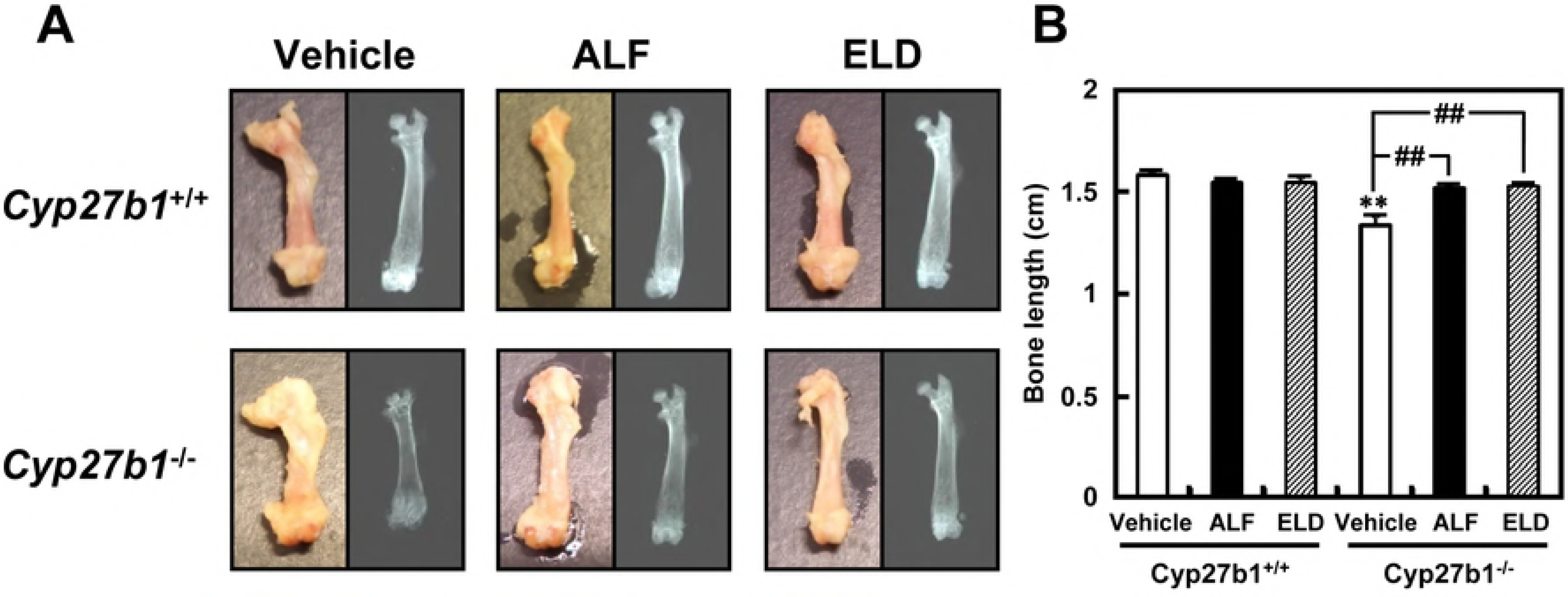
Appearance and soft X-ray image of the femurs of *Cyp27b1*^+/+^ or *Cyp27b1*^−/−^ mice administered ALF or ELD. (A) Appearance and soft X-ray image of femur. (B) Femur bone length. \*\**P* < 0.01 vs. *Cyp27b1*^+/+^ mice administered vehicle, Student’s *t*-test. ^##^P < 0.01 vs. *Cyp27b1*^−/−^ mice administered vehicle, Dunnett’s test.

### Histological staining of the femurs of *Cyp27b1*-knockout mice administered ALF or ELD

Femur calcification in *Cyp27b1*^+/+^ and *Cyp27b1*^−/−^ mice was evaluated using von Kossa staining. Areas stained black are calcification sites in which Ca has been deposited. In *Cyp27b1*^−/−^ mice administered vehicle, there were less Ca sedimentation sites in both cortical and cancellous bone than in *Cyp27b1*^+/+^ mice, suggesting hypocalcification. Bone calcification to the same degree as that seen in *Cyp27b1*^+/+^ mice was observed in *Cyp27b1*^−/−^ mice administered ALF or ELD (Fig 3A). Cartilage formation was evaluated in *Cyp27b1*^+/+^ and *Cyp27b1*^−/−^ mice by using toluidine blue staining. Areas stained purple are cartilage cells. In *Cyp27b1*^−/−^ mice administered vehicle, cartilage in the epiphyseal region and growth plate was markedly increased compared to *Cyp27b1*^+/+^ mice administered vehicle, with hypertrophy and irregularity noted in the growth-plate cartilage layer. The hypertrophy and irregularity in the growth-plate cartilage area were completely resolved by ALF or ELD administration (Fig 3B). Villanueva staining was used to evaluate osteoid formation in *Cyp27b1*^+/+^ and*Cyp27b1*^−/−^ mice. Areas stained reddish purple are osteoid. No differences in osteoid quantity were noted between *Cyp27b1*^+/+^ mice treated with vehicle, ALF, or ELD. However, osteoid tended to be present in a larger amount in *Cyp27b1*^−/−^ mice administered vehicle. In contrast, osteoid quantity decreased to the level observed in *Cyp27b1*^+/+^ mice upon administration of ALF or ELD (Fig 3C). We evaluated bone formation speed in femurs of *Cyp27b1*^+/+^ and *Cyp27b1*^−/−^ mice by double calcein labeling. Double calcein labeling was clearly observed in *Cyp27b1*^+/+^ mice administered vehicle, ALF, or ELD, indicating that bone formation was occurring normally. In contrast, calcein staining spread throughout the entire bone in *Cyp27b1*^−/−^ administered vehicle, demonstrating that bone formation did not occur at a normal speed. This abnormal bone formation speed was improved to the level in *Cyp27b1*^+/+^ mice in the ALF or ELD administration groups of *Cyp27b1*^−/−^ mice (Fig 3D).

**Fig. 3.**
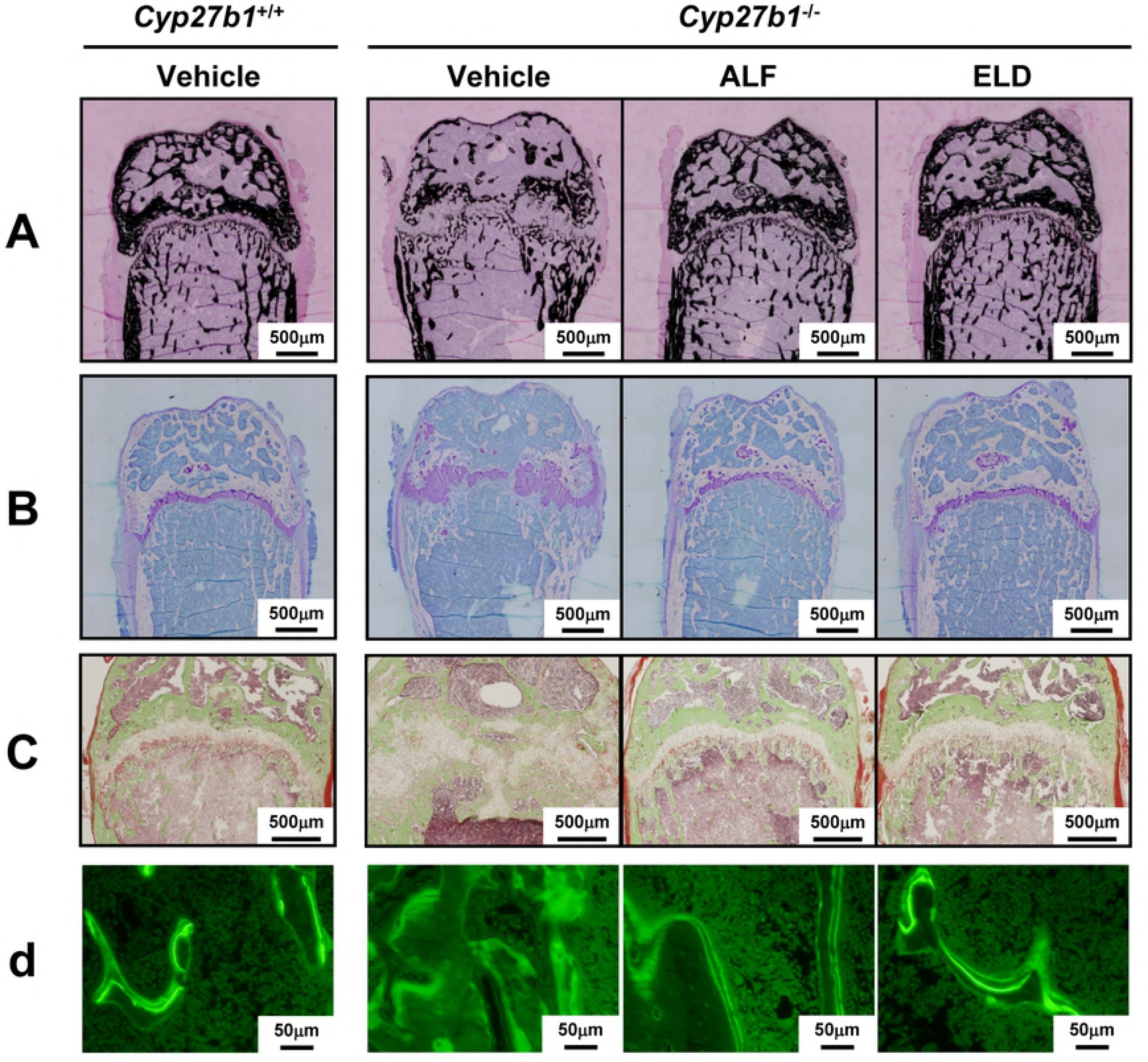
Bone histology in *Cyp27b1*^+/+^ or *Cyp27b1*^−/−^ mice administered ALF or ELD. (A) Analysis of bone mineralization by von Kossa staining. Areas stained black indicate calcification sites in which Ca has been deposited. Scale bar: 500 ゼm. (B) Analysis of cartilage formation by toluidine blue staining. Areas stained purple indicate cartilage cells. Scale bar: 500 μm. (C) Analysis of osteoid formation by Villanueva staining. Areas stained reddish purple indicate osteoid. Scale bar: 500 μm. (D) Analysis of bone formation rate by calcein double labeling. Scale bar: 50 μm.

### Femur morphology in *Cyp27b1*-knockout mice administered ALF or ELD

Femur bone density in *Cyp27b1*^+/+^ and *Cyp27b1*^−/−^ mice was measured using pQCT. In both *Cyp27b1*^+/+^ and *Cyp27b1*^−/−^ mice, overall bone density in the femoral metaphysis was significantly increased by ELD administration (Fig 4A). Although cortical bone density was significantly lower in *Cyp27b1*^−/−^ mice administered vehicle than in *Cyp27b1*^+/+^ mice administered vehicle, it increased significantly in the ALF and ELD administration groups, to the levels observed in *Cyp27b1*^+/+^ mice (Fig 4B). Cancellous bone density increased significantly in both *Cyp27b1*^+/+^ and *Cyp27b1*^−/−^ mice by administration of ELD and ALF. (Fig 4C). Femoral diaphysis cortical bone density and thickness were significantly lower in *Cyp27b1*^−/−^ than in *Cyp27b1*^+/+^ mice administered vehicle, but they increased to the level observed in *Cyp27b1*^+/+^ mice upon administration of ALF or ELD. In *Cyp27b1*^−/−^ mice administered vehicle, although horizontal bone strength, vertical bone strength, and twisting bone strength were significantly lower than in *Cyp27b1*^+/+^ mice, bone strength increased to approximately the same level as that in *Cyp27b1*^+/+^ mice in the ALF and ELD administration groups (Fig 4D, E, F). Bone strength increased particularly upon ELD administration.

**Fig. 4.**
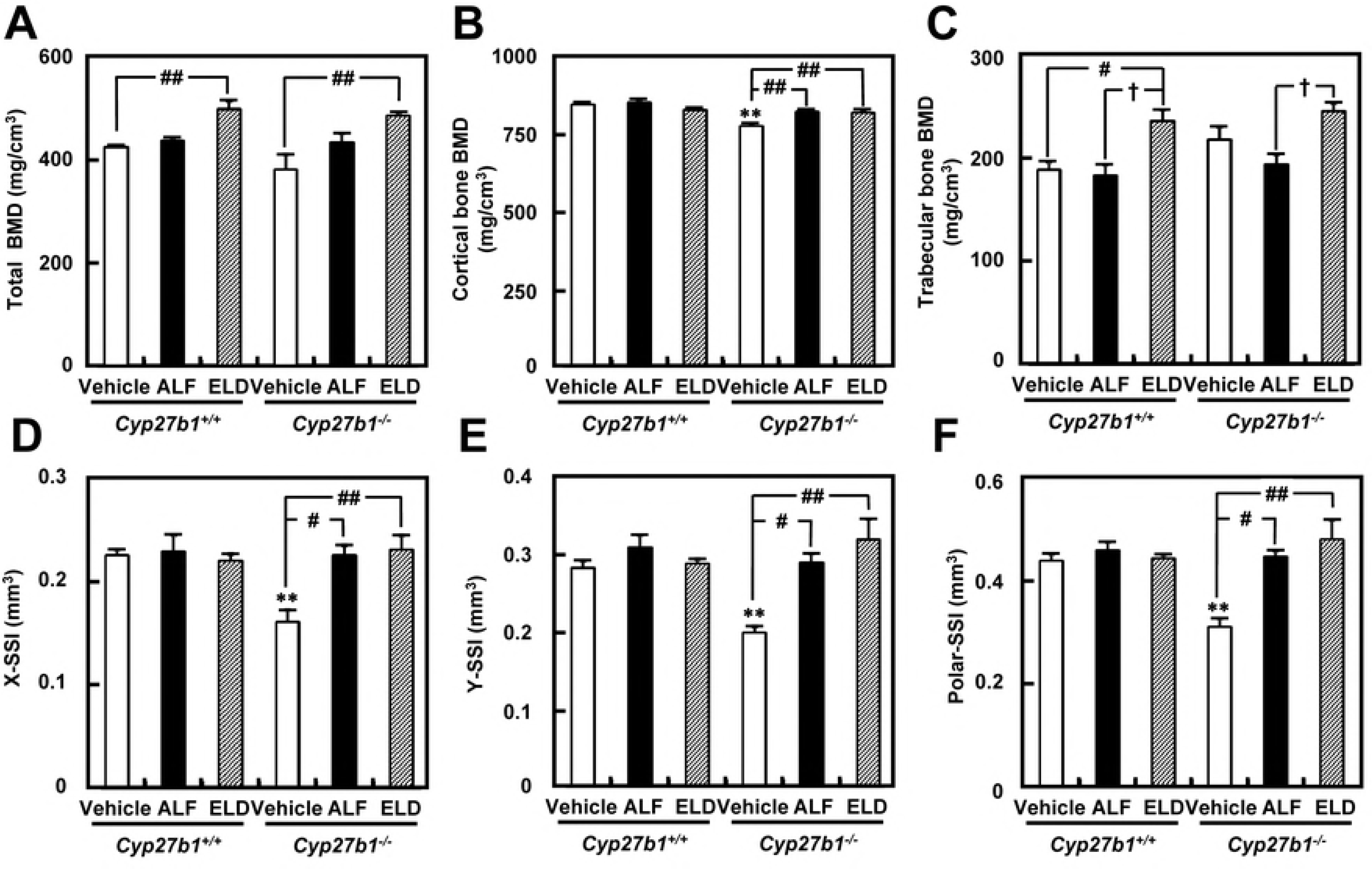
Bone mineral density and bone strength of the femurs in *Cyp27b1*^+/+^ or *Cyp27b1*^−/−^ mice administered ALF or ELD. (A) Total bone mineral density of femoral metaphysis. ##P < 0.01 vs. *Cyp27b1*^+/+^ or *Cyp27b1*^−/−^ mice administered vehicle, Student’s *t*-test. (B) Cortical bone mineral density of femoral metaphysis. \*\**P* < 0.01 vs. *Cyp27b1*^+/+^ mice administered vehicle, Student’s *t*-test. ^##^*P* < 0.01 vs. *Cyp27b1*^−/−^ mice administered vehicle, Dunnett’s test. (C) Trabecular bone density of metaphysis of the femur. ^#^*P* < 0.05 vs. *Cyp27b1*^+/+^ mice administered vehicle, Student’s *t*-test. †*P* < 0.05 vs. *Cyp27b1*^+/+^ or *Cyp27b1*^−/−^ mice administered ALF, Student’s *t*-test. (D) Bone strength in the X-axis direction of the femur. \*\**P* < 0.01 vs. *Cyp27b1*^+/+^ mice administered vehicle, Student’s *t*-test. ^#^*P* < 0.05, ^##^*P* < 0.01 vs. *Cyp27b1*^−/−^ mice administered vehicle, Dunnett’s test. (E) Bone strength in the Y-axis direction of the femur. \*\**P* < 0.01 vs. *Cyp27b1*^+/+^ mice administered vehicle, Student’s *t*-test. ^#^*P* < 0.05, ^##^*P* < 0.01 vs. *Cyp27b1*^−/−^ mice administered vehicle, Dunnett’s test. (F) Bone strength of twisted femur. \*\**P* < 0.01 vs. *Cyp27b1*^+/+^ mice administered vehicle, Student’s t-test. ^#^*P* < 0.05, ^##^*P* < 0.01 vs. *Cyp27b1*^−/−^ mice administered vehicle, Dunnett’s test.

### Structural analysis of femoral trabeculae using micro-CT in *Cyp27b1*-knockout mice administered ALF or ELD

2D micro-CT image analysis revealed a lack of trabecular bone formation in the *Cyp27b1*^−/−^ mice administered vehicle, with a markedly small amount of calcified bone, which appeared white on micro-CT images. In contrast, ALF or ELD administration restored trabecular bone formation to the level observed in *Cyp27b1*^+/+^ mice. In particular, cancellous bone formation was stronger in both *Cyp27b1*^+/+^ and *Cyp27b1*^−/−^ mice administered ELD than in *Cyp27b1*^+/+^ mice administered vehicle (Fig 5A). 3D image analysis revealed poor cancellous bone formation and hypocalcification in *Cyp27b1*^−/−^ mice administered vehicle. In contrast, trabecular bone formation and calcification were at the levels as observed in *Cyp27b1*^+/+^ mice in *Cyp27b1*^−/−^ mice treated with ALF or ELD. In particular, the ELD administration group of not only *Cyp27b1^−/−^*, but also *Cyp27b1*^+/+^ mice exhibited clearly enhanced trabecular bone formation (Fig 5B).

**Fig. 5.**
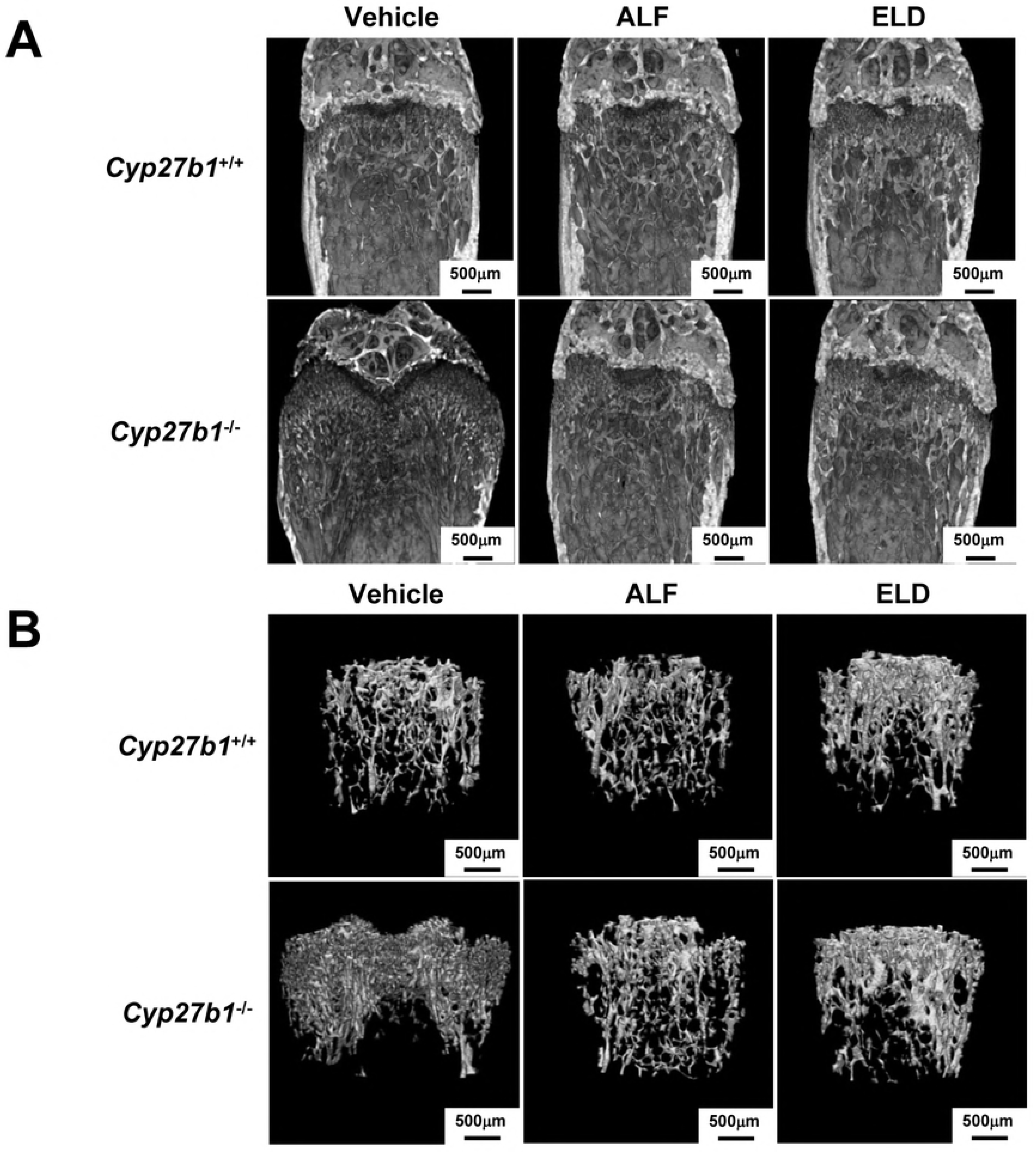
Structural analysis of trabecular bone of femur of *Cyp27b1*^+/+^ or *Cyp27b1*^−/−^ mice administered ALF and ELD. (A) 2D trabecular structure of femur by micro-CT. Scale bar: 500 μm. (B) 3D trabecular structure of femur by micro-CT. Scale bar: 500 μm.

### ALF or ELD inhibit the differentiation of ATDC5 cells

During osteogenesis, chondrocytes are formed from undifferentiated mesenchymal cells by endochondral ossification. The undifferentiated mesenchymal cells proliferate and aggregate and then differentiate into quiescent cartilage cells, proliferative cartilage cells, and hypertrophic chondrocytes, forming a cartilage layer. It is not clear which vitamin D derivative plays a role in cartilage layer formation. Therefore, we investigated the actions of vitamin D derivatives on chondrocyte differentiation using ATDC5 cells, which are capable of reproducing the chondrocyte differentiation process. Dose-dependent inhibition of cell proliferation was observed with vitamin D derivatives (Fig 6A). As a result of evaluating glycosaminoglycan expressing on chondrocytes on chondrocyte differentiation by Alcian blue staining, in vitamin D homologs and derivatives, inhibition of chondrocyte differentiation was observed in a dose-dependent manner (Fig 6B). We found that vitamin D congeners and derivatives inhibited mesenchymal cell differentiation into chondrocytes.

**Fig. 6.**
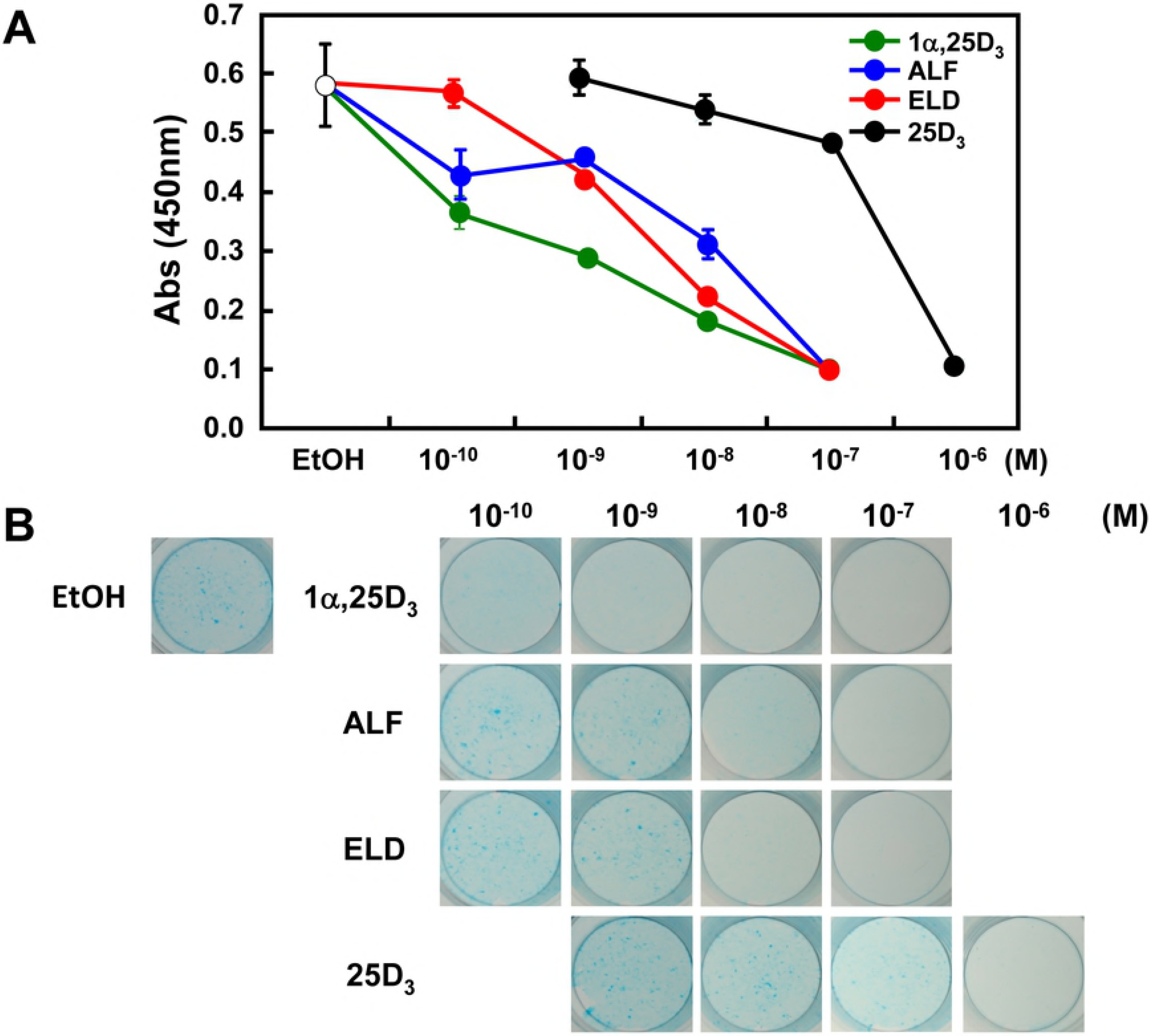
Effect of vitamin D derivatives on the proliferation and the differentiation of ATDC5 cells. (A) Proliferation of ATDC5 cells. Vitamin D homologs and vitamin D derivatives inhibited cell proliferation in a dose-dependent manner. (B) Evaluation of chondrocyte differentiation by Alcian blue staining. Vitamin D homologs as well as derivatives were found to inhibit the differentiation of cartilage cells in a dose-dependent manner.

### Effects of ALF or ELD on VDR-dependent transcriptional activity in ATDC5 cells

ATDC5 cells were used to evaluate the effects of the vitamin D derivatives on CYP24A1 transcriptional activity. 1α,25D_3_, used as a control, increased the transcriptional activity of CYP24A1 in a concentration-dependent manner. Transcriptional activity was not observed upon treatment with ALF and 25D3, but marked transcriptional activity was induced by ELD at 10^−8^ M (Fig 7A). Next, the presence or absence of DBP on VDR mediated transcriptional activity was evaluated. The effect of ALF was not influenced by DBP, but 1α,25D_3_ increased VDR-mediated transcription activity, depending on the concentration of DBP. However, the transcriptional activity induced by ELD and 25D3 decreased depending on the concentration of DBP (Fig 7B). Since ELD and 25D_3_ bind strongly to DBP, we evaluated the involvement of megalin, an uptake pathway of 25D_3_ bound to DBP. In cells in which megalin was knocked down by approximately 90% through RNAi (Fig 7C), the effects of 1α,25D_3_ and ALF were not affected, while VDR-mediated transcription activity induced by ELD and 25D3 was significantly decreased (Fig 7D).

**Fig. 7.**
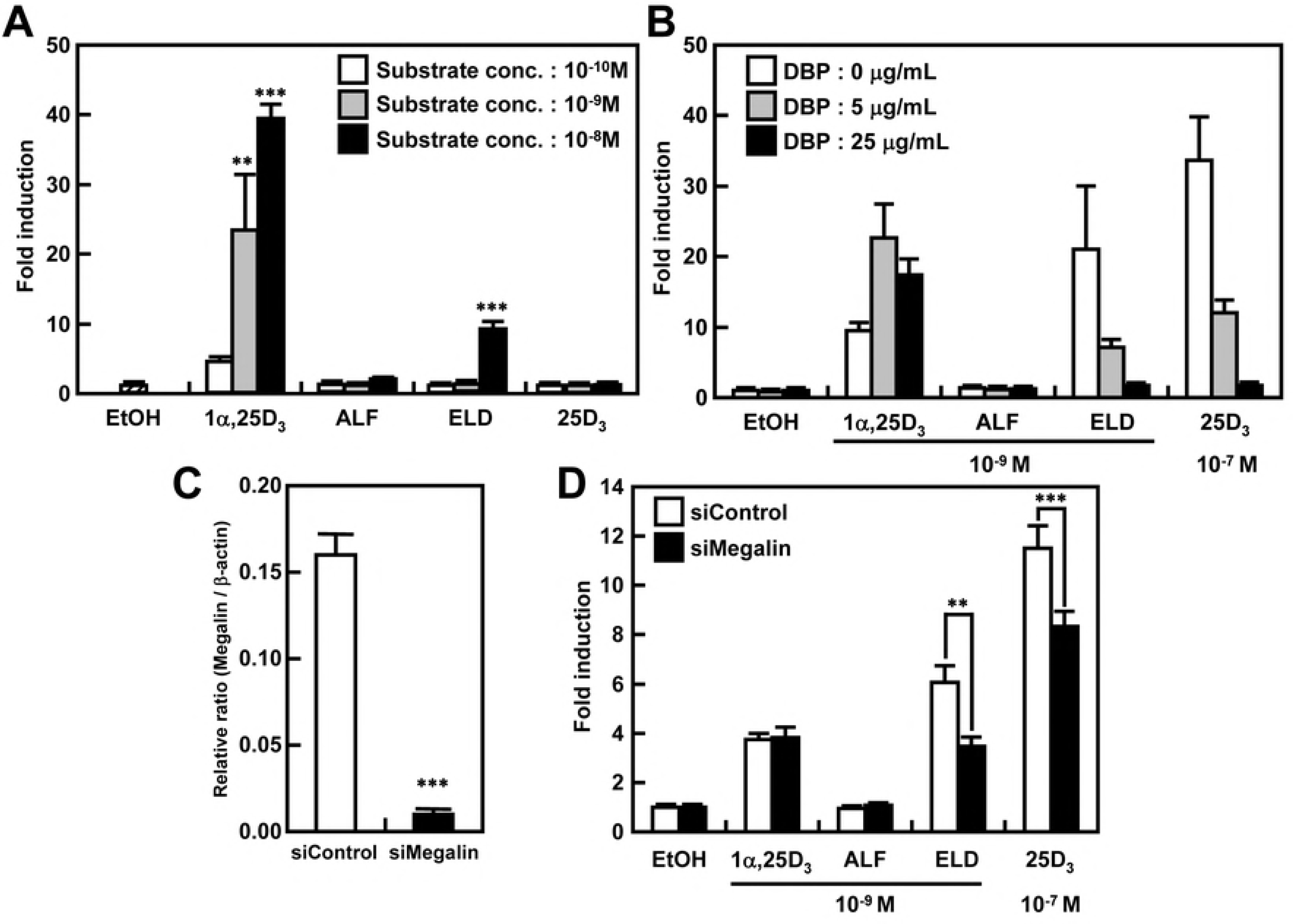
VDR-dependent transcriptional activity of vitamin D derivatives in ATDC5 cells and analysis of the effect of binding to DBP. (A) Concentration dependence of the effects of vitamin D derivatives on VDR-dependent transcriptional activity of CYP24A1 in ATDC5 cells. When 1α,25D_3_ was added, CYP24A1 transcriptional activity increased in a concentration-dependent manner. Although no transcriptional activity was noted upon treatment with ALF and 25D3, ELD markedly induced transcriptional activity at 10^−8^ M. (B) Effects of vitamin D derivatives on VDR-dependent transcriptional activity of CYP24A1 in the presence or absence of DBP in ATDC5 cells. Although ALF was not affected by DBP, 1α,25D_3_ increased CYP24A1 transcriptional activity in the presence of DBP. However, ELD and 25D_3_ showed decreased transcriptional activity in a dose-dependent manner in the presence of DBP. (C, D) Megalin-mediated VDR-dependent transcriptional activity of CYP24A1 mediated by vitamin D derivatives in the presence of DBP in ATDC5 cells. (C) Megalin mRNA expression in si-megalin-transfected ATDC5 cells. In si-megalin transfected ATDC5 cells, megalin mRNA expression was knocked down by approximately 90%. (D) VDR-dependent transcriptional activity of CYP24A1 in megalin-knockdown ATDC5 cells. In megalin-knockdown ATDC5 cells, while 1α,25D_3_ and ALF had no effect, ELD and 25D_3_ significantly lowered CYP24A1 transcriptional activity.

## Discussion

We used *Cyp27b1*^−/−^ mice, an effective model for analyzing the action of vitamin D derivatives without any influence of endogenous 1α,25D_3_, to investigate the effects of ALF and ELD administration on vitamin D metabolism. First, we evaluated the effects of ALF or ELD administration on the delayed growth observed in *Cyp27b1*^−/−^ mice. The administration of ALF or ELD restored the body weight of *Cyp27b1*^−/−^ mice to approximately the level in *Cyp27b1*^+/+^ mice. The hypocalcemia observed in *Cyp27b1*^−/−^ mice was also improved and plasma calcium was maintained at a normal level by ALF or ELD administration. In both *Cyp27b1^+/+^* and *Cyp27b1*^−/−^ mice, the plasma Ca concentration in the ELD-treated groups was relatively high, but within the normal range. In *Cyp27b1*^−/−^ mice, 1α,25D_3_ deficiency and hypocalcemia caused secondary hyperparathyroidism, with plasma PTH reaching markedly high levels [23,26]. When ALF and ELD were administered, plasma PTH decreased markedly. In a previous study, when 1α,25D_3_ was administered to *Cyp27b1*^−/−^ mice, hyperparathyroidism was improved [27]. ALF and ELD both correct hyperparathyroidism via the same process [27]. By promoting active Ca absorption in the intestinal tract and kidneys, 1α,25D_3_ maintains Ca homeostasis in the plasma. Active Ca absorption occurs when Ca flows in through the epithelial Ca channel ECaC2, after which Ca is transported into cells by proteins such as calbindin-D9k. The VDRE is found upstream of vitamin D-response gene promoters, and its expression is controlled at the transcriptional level by 1α,25D_3_ [28,29]. Therefore, we measured the mRNA expression levels of *ECaC2* and *calbindin-D_9k_* in the duodenum to further investigate the Ca metabolism-modulating effects of ALF and ELD. In accordance with a previous report [30], administering ALF to mice did not induce *ECaC2* or *calbindin-D*_*9k*_ expression in the intestinal tract. However, ELD increased the mRNA expression of *ECaC2* and *calbindin-D*_*9k*_ in both *Cyp27b1*^+/+^ and *Cyp27b1*^−/−^ mice, most likely because ELD is an active ligand that binds directly with VDR. Even in experiments using ATDC5 cells, ELD increased VDR-mediated transcriptional activity in a concentration-dependent manner, similar to 1α,25D_3_. (Fig 7A). These results demonstrated that the administration of ALF or ELD made it possible to maintain the plasma Ca in *Cyp27b1*^−/−^ mice within the normal range, because it enhanced active Ca absorption mediated by ECaC2 and calbindin-D9k expression in the duodenum.

It has been reported that ELD reduces plasma 1α,25D_3_ and increases plasma 24,25D_3_ in experimental animals [22]. This appears to be because ELD induces CYP24A1, which metabolizes 1α,25D_3_ and 25D_3_ into inactive forms. Thus, lowering the plasma 1α,25D_3_ concentration appears to be one of the physiological mechanisms of ELD. When we investigated the potential influence of ALF and ELD on mRNA expression of *CYP2R1* and *CYP27A1* in the liver, no effects were noted. Because studies using *Cyp27b1*^−/−^ mice raised on high-calcium feed and vdr-knockout mice revealed that the expression of CYP2R1 and CYP27A1 is similar to that in *Cyp27b1*^+/+^ mice [26], it appears that 1α,25D_3_ and 25D_3_ do not control CYP2R1 and CYP27A1 expression. The mRNA expression of *CYP24A1* in the kidneys of *Cyp27b1*^−/−^ mice was significantly lower than that in *Cyp27b1*^+/+^ mice. However, *CYP24A1* expression was strongly induced by ALF and ELD administration. Similar effects were noted in *Cyp27b1*^+/+^ mice. Because VDRE is present upstream from the *CYP24A1* promoter [31], it appears that, as with 1α,25D_3_, ELD interacts with the VDR to induce *CYP24A1* expression. Moreover, because of its high DBP bindability, ELD is stable in the plasma and circulates longer than 1α,25D_3_ [32]. Therefore, it appears that ELD more strongly induces *CYP24A1* expression than ALF. Like ELD, it has been reported that the compound 2α-hydroxypropoxy-1α,25D_3_ (O2C3) having a 3-hydroxypropoxy group at the 2α position is more difficult to metabolize at CYP24A1 than 1α,25D_3_ [33], which may have allowed for the prolonged effects of ELD. Based on the above, the significant effects of ELD that were noted even in *Cyp27b1*^+/+^ mice indicated that ELD can promote Ca absorption and improve osteogenesis in patients with osteoporosis. By using *Cyp27b1*^−/−^ mice in our experiment, we were able to demonstrate that ELD exhibits Ca metabolism-regulating effects without being influenced by endogenous 1α,25D_3_.

A study in OVX rats showed that ELD is superior to 1α,25D_3_ in terms of increasing lumbar vertebrae and femur bone density, and inhibiting bone resorption [24]. However, ELD administration decreases the plasma 1α,25D_3_ concentration [25], and this decrease may mediate the action of ELD on bones. Previous studies using OVX rats were unable to exclude the influence of 1α,25D_3_ to analyze the actions of ELD alone. Therefore, in order to clarify the actions of ELD alone on bone, excluding the effects of 1α,25D_3_, we analyzed the osteogenic actions of ALF and ELD in *Cyp27b1*^−/−^ mice. Externally, the femurs and tibias in vehicle-treated *Cyp27b1*^−/−^ mice were observed to have hypertrophy in the epiphyseal region and reduced bone length. However, the administration of ALF or ELD improved these abnormalities to a state similar to that observed in *Cyp27b1*^+/+^ mice. Bone samples of *Cyp27b1*^−/−^ mice exhibited hypocalcification, hypertrophy and irregularity in the growth-plate cartilage, and increased osteoid formation, as well as poor osteogenesis speed and clear lesions, indicating imperfect osteogenesis. These abnormalities were completely restored by ALF and ELD administration, and osteogenesis occurred at the level observed in *Cyp27b1^+I+^* mice. Osteogenic abnormalities involving bone length and bone density in *Cyp27b1*^−/−^ mice can be sufficiently improved with Ca supplementation [26]. However, hypocalcification and growth-plate cartilage hypertrophy cannot be completely normalized with Ca supplementation alone. Accordingly, the fact that osteogenesis occurred at the same level as in *Cyp27b1*^+/+^ mice due to ALF or ELD administration indicates that not only Ca metabolism was maintained, but 1α,25D_3_ was metabolized and generated from ALF as well as the direct actions of ELD on the bone. Previously, it has been reported that when ATDC5 cells, which can differentiate into cartilage cells, are cultured to differentiate into cartilage cells, the addition of 1α,25D_3_ markedly inhibits such differentiation [34]. We found that when ATDC5 cells were treated with not only 1α,25D_3_, but also ALF and ELD, ATDC5 cell proliferation and differentiation were inhibited in a dose-dependent manner (Fig 6). We also investigated whether the presence or absence of DBP affects VDR transcriptional activity. The results showed that while the effect of ALF was not at all affected by DBP, 1α,25D_3_ increased VDR transcriptional activity when DBP was present. However, ELD and 25D3 decreased VDR-mediated transcriptional activity in a concentration-dependent manner (Fig 7B). Because ELD and 25D_3_ are thought to strongly bind to DBP, we evaluated the involvement of megalin, an uptake route for 25D_3_ bound to DBP. Cells in which megalin had been knocked down to approximately 90% using RNAi were found to not be affected by 1α,25D_3_ or ALF. However, VDR-mediated transcriptional activity significantly decreased for ELD and 25D_3_ (Fig 7D). Based on the above results, it appears that ELD strongly binds with DBP to act on cartilage cells mediating megalin so as to maintain their normal proliferation and differentiation.

When we used non-decalcified specimens to measure parameters related to bone structure, osteogenesis, and bone resorption, we found that *Cyp27b1*^−/−^ vehicle-treated mice exhibited decreased bone quantity, decreased trabecula width and number, abnormally widened trabecular spaces, and decreased calcification and osteogenesis. However, ALF or ELD administration improved these abnormalities to the normal levels of these parameters measured in *Cyp27b1*^+/+^ mice (S1 Table). In the *Cyp27b1*^−/−^ ELD administration group, in particular, bone resorption surface, osteoclast count, and osteoclast surface tended to decrease more than during ALF administration, and it was found that bone quantity and trabeculae were significantly increased, while the osteogenic rate was maintained. This suggests that ELD more strongly inhibits bone resorption than ALF, resulting in superior osteogenesis. Similarly, in a previous, the femurs of mice administered ELD exhibited decreased osteoclast counts [35]. Moreover, as the lumbar vertebrae of OVX rats administered ELD were reported to exhibit greater decreases in osteoclast surface than those administered ALF, ELD appears to have strong bone resorption-inhibitory effects [36].

ELD was found to promote femur cancellous bone density in both *Cyp27b1*^+/+^ and *Cyp27b1*^−/−^ mice more strongly than ALF. Moreover, the fact that the density-increasing action of ELD on cancellous bone was also observed in *Cyp27b1*^+/+^ mice demonstrated that ELD exhibits particularly strong effects on cancellous bone. This may be due to differences in bone turnover between cortical and cancellous bone. Although no remodeling occurs in the cortical bone of mice and rats, bone resorption by osteoclasts and osteogenesis by osteoblasts occurs on the trabecular surface of cancellous bone, implying that remodeling occurs continuously [37]. This suggests that ELD directly acts on the osteoclasts and osteoblasts in cancellous bone, where more bone remodeling occurs than in cortical bone, to reduce the osteoclast count and inhibit bone resorption, thereby promoting osteogenesis by osteoblasts. The particularly strong effects of ELD compared to ALF may have been due to ELD not only having high bone selectivity [21] (Fig 7A), but also higher DBP-bindability than ALF and a long plasma half-life [16–19]. Previously, it has been reported that 1α,25D_3_ markedly inhibits c-Fos expression in osteoclast precursor cells, thereby suppressing osteoclast differentiation [38]. Therefore, ELD may also directly act on osteoclast precursor cells via this mechanism to inhibit bone resorption [39]. Accordingly, the direct action of ELD on osteoblasts is likely to have induced the expression of factors that promote osteogenesis, such as osteocalcin (Fig 8). In this study, we have not evaluated the effect of ELD on mini-modeling, a bone formation without prior bone resorption that has been shown in rat trabecular bone treated with ELD [40]. By using *Cyp27b1*^−/−^ mice, it seems that the osteogenic effect of ELD on mini-modeling could be analyzed in more detail.

**Fig. 8.**
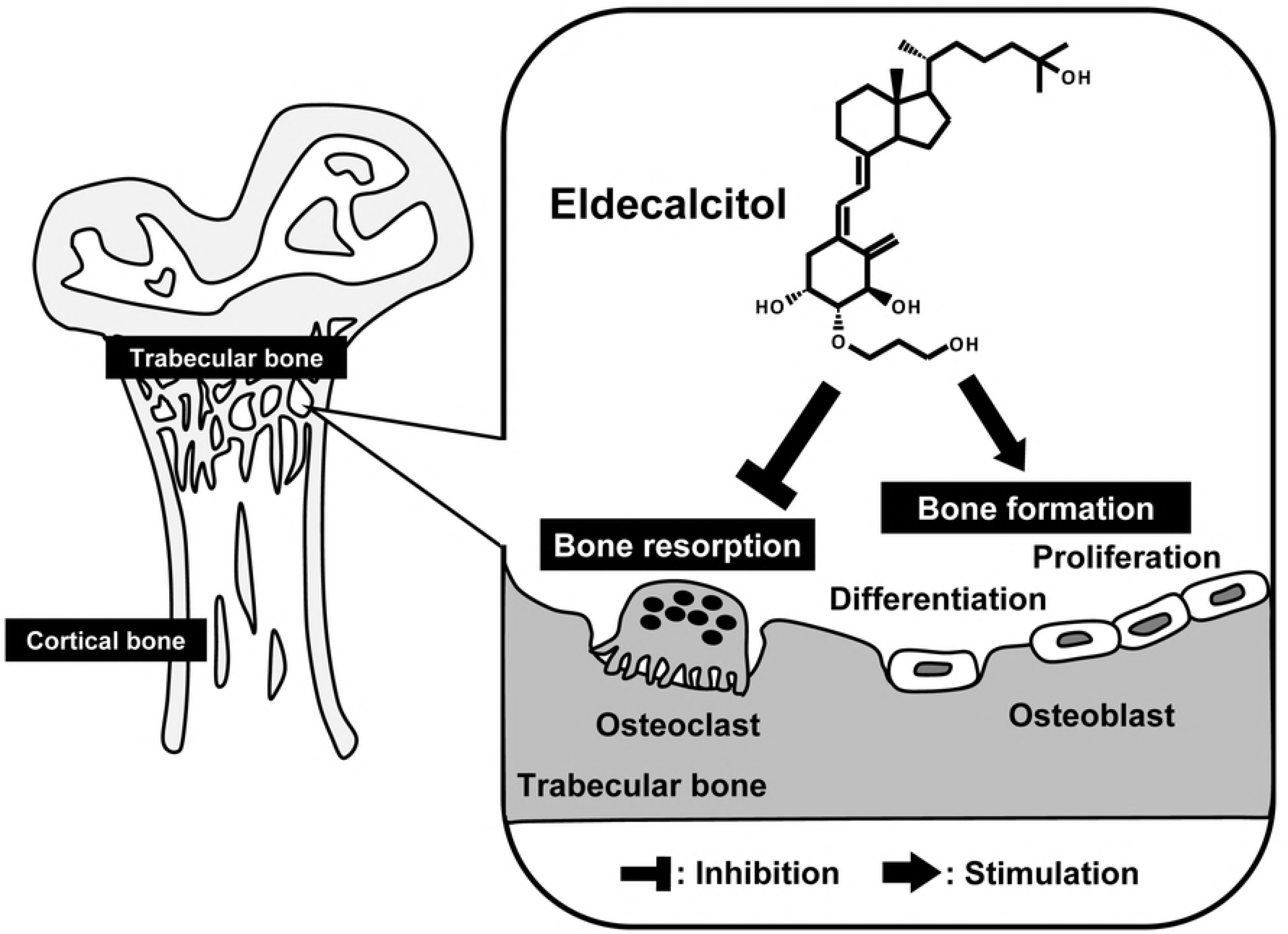
Schematic illustration of ELD action in trabecular bone.

Although bone strength was markedly decreased in *Cyp27b1*^−/−^ mice treated with vehicle, the administration of ALF and ELD allowed these animals to maintain bone strength at a level similar to that of *Cyp27b1*^+/+^ mice (Fig 4). In the *Cyp27b1*^−/−^ ELD administration group, in particular, bone strength tended to be more strongly increased than in the ALF administration group. In terms of factors related to bone strength, the trabecular structure is important in addition to bone density. Thinner trabeculae in cancellous bone result in more fragile bones. Using 2D and 3D micro-CT analyses of femoral trabeculae, we found that trabecular bone formation and calcification were markedly decreased in *Cyp27b1*^−/−^ mice administered vehicle. However, both were maintained at the same level as in *Cyp27b1*^+/+^ mice upon ALF and ELD administration. In the ELD administration group in particular, trabecular bone formation was clearly enhanced in not only *Cyp27b1*^−/−^ mice, but in also *Cyp27b1*^+/+^ mice, demonstrating that ELD strongly increases trabecular formation and trabecular connectivity. These results suggest that ELD acts more strongly on cancellous bone, in which remodeling occurs and turnover is faster than in cortical bone, to promote osteogenesis and inhibit bone resorption. Osteoporosis, which commonly occurs in postmenopausal women, develops as decreases in female hormones that act to inhibit bone resorption and cause bone resorption to become relatively dominant. Because postmenopausal osteoporosis involves particularly marked decreases in the trabecular count in cancellous bone [41], ELD may be a very effective therapeutic agent for the disease due to this selective action on cancellous bone and promotion of bone density and trabecular bone formation. We administered ELD to weaned mice and evaluated the effects on osteogenesis, revealing that it exhibits highly superior osteogenesis effects. This indicates that ELD is also an effective agent for treating vitamin D-dependent rickets type I.

## Acknowledgements

We thank Dr. Keiichi Ozono for providing pGVB2-rat Cyp24a1 vector, pM-human VDR and pGVP2-GAL2. We thank Chugai Pharmaceutical Co., Ltd. for providing medium-chain triglycerides, ALF, and ELD. We thank numerous members of the Okano laboratories for helpful technical assistance and discussions on the manuscript.

## Author contributions

Conceived and designed the experiments: YH, KN, TO. Performed the experiments: YH, KN, KI, MK. Analyzed the data: YH, KN, KI. Contributed reagents/materials/analysis tools: NK, YS. Wrote the paper: YH, TS, TO. Discussed the paper: YH, KN, TS, NO, TO.

## Supporting information

**Table S1. Measurement of bone morphology of *Cyp27b1*^+/+^ or *Cyp27b1*^−/−^ mice administered ALF or ELD for 3 or 6 weeks.**

\**P* < 0.05 vs. *Cyp27b1*^+/+^ mice administered vehicle, Student’s *t*-test. ^#^*P* < 0.05 vs. *Cyp27b1*^−/−^ mice administered vehicle, Dunnett’s test.

## References

1. Daiger SP, Schanfield MS, Cavalli-Sforza LL. Group-specific component (Gc) proteins bind vitamin D and 25-hydroxyvitamin D. Proc Natl Acad Sci U S A. 1975;72:2076–2080.

2. Baker AR, McDonnell DP, Hughes M, Crisp TM, Mangelsdorf DJ, Haussler MR, Pike JW, Shine J, O’Malley BW. Cloning and expression of full-length cDNA encoding human vitamin D receptor. Proc Natl Acad Sci USA. 1988;85:3294–3298.

3. Kliewer SA, Umesono K, Mangelsdorf DJ, Evans RM. Retinoid X receptor interacts with nuclear receptors in retinoic acid, thyroid hormone and vitamin D3 signalling. Nature. 1992;355:446–449. DOI:10.1038/355446a0

4. Rachez C, Freedman LP. Mechanisms of gene regulation by vitamin D_3_ receptor: a network of coactivator interactions. Gene, 2000;246:9–21.

5. Darwish H, DeLuca HF. Vitamin D-regulated gene expression. Crit. Rev. Eukaryot. Gene Exp. 1993;3:89–116.

6. Shams-White MM, Chung M1, Du M, Fu Z, Insogna KL, Karlsen MC, LeBoff MS, Shapses SA, Sackey J, Wallace TC, Weaver CM. Dietary protein and bone health: a systematic review and meta-analysis from the National Osteoporosis Foundation. Am J Clin Nutr. 2017;105:1528–1543. DOI:10.3945/ajcn.116.145110

7. Galante L, Colston KW, MacAuley SJ, MacIntyre I. Effect of calcitonin on vitamin D metabolism. Nature. 1972;238:271–273.

8. Takeyama K, Kitanaka S, Sato T, Kobori M, Yanagisawa J, Kato S. 25-Hydroxyvitamin D3 1alpha-hydroxylase and vitamin D synthesis. Science. 1997;277:1827–1830.

9. Sairanen S, Kärkkäinen M, Tähtelä R, Laitinen K, Mäkelä P, Lamberg-Allardt C, Välimäki MJ. Bone mass and markers of bone and calcium metabolism in postmenopausal women treated with 1,25-dihydroxyvitamin D (Calcitriol) for four years. Calcif Tissue Int. 2000;67:122–127.

10. Murayama E, Miyamoto K, Kubodera N, Mori T, Matsunaga I. Synthetic studies of vitamin D_3_ analogues. VIII. Synthesis of 22-oxavitamin D_3_ analogues. Chem Pharm Bull. 1986;34:4410–4413.

11. Hayashi Y., Fujita T., Inoue T. Decrease of vertebral fracture in osteoporotics by administration of 1α-hydroxy-vitamin D_3_. J Bone Mineral Metab. 1992;10:184–188. DOI: https://doi.org/10.1007/BF02378983

12. Orimo H, Shiraki M, Hayashi Y, Hoshino T, Onaya T, Miyazaki S, Kurosawa H, Nakamura T, Ogawa N. Effects of 1 alpha-hydroxyvitamin D3 on lumbar bone mineral density and vertebral fractures in patients with postmenopausal osteoporosis. Calcif Tissue Int. 1994;54:370–376.

13. Shiraishi A, Higashi S, Masaki T, Saito M, Ito M, Ikeda S, Nakamura T. A comparison of alfacalcidol and menatetrenone for the treatment of bone loss in an ovariectomized rat model of osteoporosis. Calcif Tissue Int. 2002;71:69–79.

14. Morimoto S, Imanaka S, Koh E, Shiraishi T, Nabata T, Kitano S, Miyashita Y, Nishii Y, Ogihara T. Comparison of the inhibitions of proliferation of normal and psoriatic fibroblasts by 1 alpha,25-dihydroxyvitamin D_3_ and synthetic analogues of vitamin D_3_ with an oxygen atom in their side chain. Biochem Int. 1989;19:1143–1149.

15. Yasuda K, Iwanaga Y, Ogawa K, Mano H, Ueno S, Kimoto S, Ohta M, Kamakura M, Ikushiro S, Sakaki T. Human hepatic metabolism of the anti-osteoporosis drug eldecalcitol involves sterol C4-methyl oxidase. Pharmacol Res Perspect. 2015;3:e00120. DOI:10.1002/prp2.120

16. Miyamoto K, Murayama E, Ochi K, Watanabe H, Kubodera N. Synthetic studies of vitamin D analogues. XIV. Synthesis and calcium regulating activity of vitamin D_3_ analogues bearing a hydroxyalkoxy group at the 2 beta-position. Chem Pharm Bull. 1993;41:1111–1113.

17. Okano T, Tsugawa N, Masuda S, Takeuchi A, Kobayashi T, Takita Y, Nishii Y. Regulatory activities of 2 beta-(3-hydroxypropoxy)-1 alpha, 25-dihydroxyvitamin D3, a novel synthetic vitamin D3 derivative, on calcium metabolism. Biochem Biophys Res Commun. 1989;163:1444–1449.

18. Serizawa K, Yogo K, Tashiro Y, Takeda S, Kawasaki R, Aizawa K, Endo K. Eldecalcitol prevents endothelial dysfunction in postmenopausal osteoporosis model rats. J Endocrinol. 2016;228:75–84. DOI:10.1530/JOE-15-0332

19. Hatakeyama S, Nagashima S, Imai N, Takahashi K, Ishihara J, Sugita A, Nihei T, Saito H, Takahashi F, Kubodera N. Synthesis and evaluation of a 3-position diastereomer of 1α,25-dihydroxy-2β-(3-hydroxypropoxy)vitamin D_3_ (ED-71). Bioorg Med Chem. 2006;14:8050–8056.

20. Tsurukami H, Nakamura T, Suzuki K, Sato K, Higuchi Y, Nishii Y. A novel synthetic vitamin D analogue, 2β-(3-hydroxypropoxy)1α, 25-dihydroxyvitamin D_3_ (ED-71), increases bone mass by stimulating the bone formation in normal and ovariectomized rats. Calcif Tissue Int. 1994;54:142–149.

21. Kubodera N, Tsuji N, Uchiyama Y, Endo K. A new active vitamin D s analog, ED-71, causes increase in bone mass with preferential effects on bone in osteoporotic patients. J Cell Biochem. 2003;88:286–289.

22. Matsumoto T, Miki T, Hagino H, Sugimoto T, Okamoto S, Hirota T, Tanigawara Y, Hayashi Y, Fukunaga M, Shiraki M, Nakamura T. A new active vitamin D, ED-71, increases bone mass in osteoporotic patients under vitamin D supplementation: a randomized, double-blind, placebo-controlled clinical trial. J Clin Endocrinol Metab. 2005;90:5031–5036.

23. Dardenne O, Prud’homme J, Arabian A, Glorieux FH, St-Arnaud R. Targeted inactivation of the 25-hydroxyvitamin D_3_-1αhydroxylase gene (CYP27B1) creates an animal model of pseudovitamin D-deficiency rickets. Endocrinology. 2001;142:3135–3141.

24. Panda DK, Miao D, Tremblay ML, Sirois J, Farookhi R, Hendy GN, Goltzman D. Targeted ablation of the 25-hydroxyvitamin D 1α-hydroxylase enzyme: evidence for skeletal, reproductive, and immune dysfunction. Proc Natl Acad Sci U S A. 2001;98:7498–7503.

25. Vanhooke JL1, Prahl JM, Kimmel-Jehan C, Mendelsohn M, Danielson EW, Healy KD, DeLuca HF. CYP27B1 null mice with LacZreporter gene display no 25-hydroxyvitamin D_3_-1α-hydroxylase promoter activity in the skin. Proc Natl Acad Sci U S A. 2006;103:75–80. DOI:10.1073/pnas.0509734103

26. Hirota Y, Nakagawa K, Mimatsu S, Sawada N, Sakaki T, Kubodera N, Kamao M, Tsugawa N, Suhara Y, Okano T. Nongenomic effects of 1α,25-dihydroxyvitamin D3 on cartilage formation deduced from comparisons between Cyp27b1 and Vdr knockout mice. Biochem Biophys Res Commun. 2017;483:359–365. DOI:10.1016/j.bbrc.2016.12.139

27. Dardenne O, Prud’homme J, Glorieux FH, St-Arnaud R. Rescue of the Pseudo - Vitamin D Deficiency Rickets Phenotype of CYP27B1 - Deficient Mice by Treatment With 1,25 - Dihydroxyvitamin D_3_: Biochemical, Histomorphometric, and Biomechanical Analyses. J Steroid Biochem Mol Biol. 2004;89-90:327–330. DOI:10.1016/j.jsbmb.2004.03.026

28. Gill RK, Christakos S. Identification of sequence elements in mouse calbindin-D28k gene that confer 1,25-dihydroxyvitamin D3- and butyrate-inducible responses. Proc Natl Acad Sci U S A. 1993;90:2984–2988.

29. Colnot S, Ovejero C, Romagnolo B, Porteu A, Lacourte P, Thomasset M, Perret C. Transgenic analysis of the response of the rat calbindin-D 9k gene to vitamin D. Endocrinology, 2000;141:2301–2308. DOI:10.1210/endo.141.7.7557

30. Ishizawa M, Matsunawa M, Adachi R, Uno S, Ikeda K, Masuno H, Shimizu M, Iwasaki K, Yamada S, Makishima M. Lithocholic acid derivatives act as selective vitamin D receptor modulators without inducing hypercalcemia. J Lipid Res. 2008;49:763–772. DOI:10.1194/jlr.M700293-JLR200

31. Ohyama Y, Noshiro M, Eggertsen G, Gotoh O, Kato Y, Björkhem I, Okuda K. Structural characterization of the gene encoding rat 25-hydroxyvitamin D3 24-hydroxylase. Biochemistry. 1993;32:76–82.

32. Sanford M, McCormack PL. Eldecalcitol: a review of its use in the treatment of osteoporosis. Drugs. 2011;71:1755–1770. DOI:10.2165/11206790-000000000-00000

33. Abe D, Sakaki T, Kusudo T, Kittaka A, Saito N, Suhara Y, Fujishima T, Takayama H, Hamamoto H, Kamakura M, Ohta M, Inouye K. Metabolism of 2α-propoxy-1α,25-dihydroxyvitamin D_3_ and 2α-(3-hydroxypropoxy)-1α, 25-dihydroxyvitamin D_3_ by human CYP27A1 and CYP24A1. Drug Metab Dispos. 2005;33,778–784. DOI:10.1124/dmd.104.003038

34. Akiyama H, Hiraki Y, Shigeno C, Kohno H, Shukunami C, Tsuboyama T, Kasai R, Suzuki F, Konishi J, Nakamura T. 1α,25-dihydroxyvitamin D_3_ inhibits cell growth and chondrogenesis of a clonal mouse EC cell line, ATDC5. J Bone Miner Res. 1996;11:22–28. DOI:10.1002/jbmr.5650110105

35. Okuda N, Takeda S, Shinomiya K, Muneta T, Itoh S, Noda M, Asou Y. ED-71, a novel vitamin D analog, promotes bone formation and angiogenesis and inhibits bone resorption after bone marrow ablation. Bone. 2007;40:281–292. DOI:10.1016/j.bone.2006.08.017

36. Uchiyama Y, Higuchi Y, Takeda S, Masaki T, Shira-Ishi A, Sato K, Kubodera N, Ikeda K, Ogata E. ED-71, a vitamin D analog, is a more potent inhibitor of bone resorption than alfacalcidol in an estrogen-deficient rat model of osteoporosis. Bone. 2002;30:582–588.

37. Parfitt AM. Misconceptions (2): turnover is always higher in cancellous than in cortical bone. Bone. 2002;30:807–9.

38. Takasu H, Sugita A, Uchiyama Y, Katagiri N, Okazaki M, Ogata E, Ikeda K. c-Fos protein as a target of anti-osteoclastogenic action of vitamin D, and synthesis of new analogs. J Clin Invest. 2006;116:528–535. DOI:10.1172/JCI24742

39. Ikeda T, Kohno H, Yamamuro T, Kasai R, Ohta S, Okumura H, Konishi J, Kikuchi H, Shigeno C. The effect of active vitamin D_3_ analogs and dexamethasone on the expression of osteocalcin gene in rat tibiae in vivo. Biochem Biophys Res Commun. 1992;189:1231–1235.

40. Saito H, Takeda S, Amizuka N. Eldecalcitol and calcitriol stimulates ‘bone minimodeling,’ focal bone formation without prior bone resorption, in rat trabecular bone. J Steroid Biochem Mol Biol. 2013;136:178–182.

41. Riggs BL, Khosla S, Melton LJ 3rd. A unitary model for involutional osteoporosis: estrogen deficiency causes both type I and type II osteoporosis in postmenopausal women and contributes to bone loss in aging men. J Bone Miner Res. 1998;13:763–73. DOI:10.1359/jbmr.1998.13.5.763

